# Stain-free nucleus identification in holographic learning flow cyto-tomography

**DOI:** 10.1101/2021.12.22.473826

**Authors:** Daniele Pirone, Joowon Lim, Francesco Merola, Lisa Miccio, Martina Mugnano, Vittorio Bianco, Flora Cimmino, Feliciano Visconte, Annalaura Montella, Mario Capasso, Achille Iolascon, Pasquale Memmolo, Demetri Psaltis, Pietro Ferraro

**Affiliations:** CNR-ISASI, Institute of Applied Sciences and Intelligent Systems “E. Caianiello”, Via Campi Flegrei 34, 80078 Pozzuoli, Napoli, Italy; DIETI, Department of Electrical Engineering and Information Technologies, University of Naples “Federico II”, via Claudio 21, 80125 Napoli, Italy; EPFL, Ecole Polytechnique Fédérale de Lausanne, Optics Laboratory, CH-1015 Lausanne, Switzerland; CEINGE - Advanced Biotechnologies, Via Gaetano Salvatore 486, 80131 Napoli, Italy; DMMBM, Department of Molecular Medicine and Medical Biotechnology, University of Naples “Federico II”, Via Pansini 5, 80131 Napoli, Italy

## Abstract

Quantitative Phase Imaging (QPI) has gained popularity because it can avoid the staining step, which in some cases is difficult or impossible. However, QPI does not provide the well-known specificity to various parts of the cell (e.g., organelles, membrane). Here we show a novel computational segmentation method based on statistical inference that bridges the gap between the specificity of Fluorescence Microscopy (FM) and the label-free property of QPI techniques to identify the cell nucleus. We demonstrate application to stain-free cells reconstructed through the holographic learning and in flow cyto-tomography modality. In particular, by means of numerical simulations and two cancer cell lines, we demonstrate that the nucleus-like regions can be accurately distinguished within the stain-free tomograms. We show that our experimental results are consistent with confocal FM data and microfluidic cytofluorimeter outputs. This is a significant step towards extracting the three-dimensional (3D) intracellular specificity directly from the phase-contrast data in a typical flow cytometry configuration.

## Introduction

Traditional tools of histopathology will evolve soon and the future of precision medicine will pass through the accurate screening of single cells. A key challenge that will allow the next jump forward is achieving a more informative label-free microscopy. Nowadays, the gold standard imaging tool in cell biology is FM, in which stains or fluorescent tags are used to make the biological sample visible on a selective basis^1,2^. However, FM is generally qualitative and limited by photo-bleaching and photo-toxicity^2^, and sample preparation protocols can be expensive, time-consuming, and operator-sensitive. Therefore, avoiding staining will permit one to access non-destructive, rapid, and chemistry-free analysis in biology and medicine. On the other hand, QPI is emerging as a very useful tool in label-free microscopy, and recently many significant results have been achieved in this field^2-11^. Starting from QPI, the Optical Diffraction Tomography (ODT) can be performed to map the refractive index (RI) of a biological specimen in 3D, thus providing a full quantitative measurement of 3D morphologies and RI volumetric distributions at the single-cell level^12-20^. However, the advantages of QPI are counterbalanced by the lack of intra-cellular specificity. Among all intra-cellular structures, the nucleus is the principal one in the eukaryotic cell since it contains most of the cellular genetic material and it is responsible for the cellular lifecycle. Identifying the nucleus-like region in label-free 3D imaging is a challenging task since the nuclear size and RI can vary among different cell lines, within the same cell line, and even within the same cell, depending on the lifecycle’s phases. In addition, different subcellular structures show similar RI values^21^, thus making any threshold-based detection method ineffective. Recently, significant progresses have been reported to introduce specificity in QPI by artificial intelligence (AI). In particular, deep learning was employed to virtually stain unlabelled tissues^22-24^ as well as single cells^25,26^ in quantitative phase maps (QPMs), and to identify nuclei of unlabelled and adhered cells in 3D ODT reconstructions^27^. To illustrate the state of the art for intra-cellular specificity, a summary diagram about the comparison between label-free and fluorescent techniques is shown in Fig. 1.

**Fig. 1.**
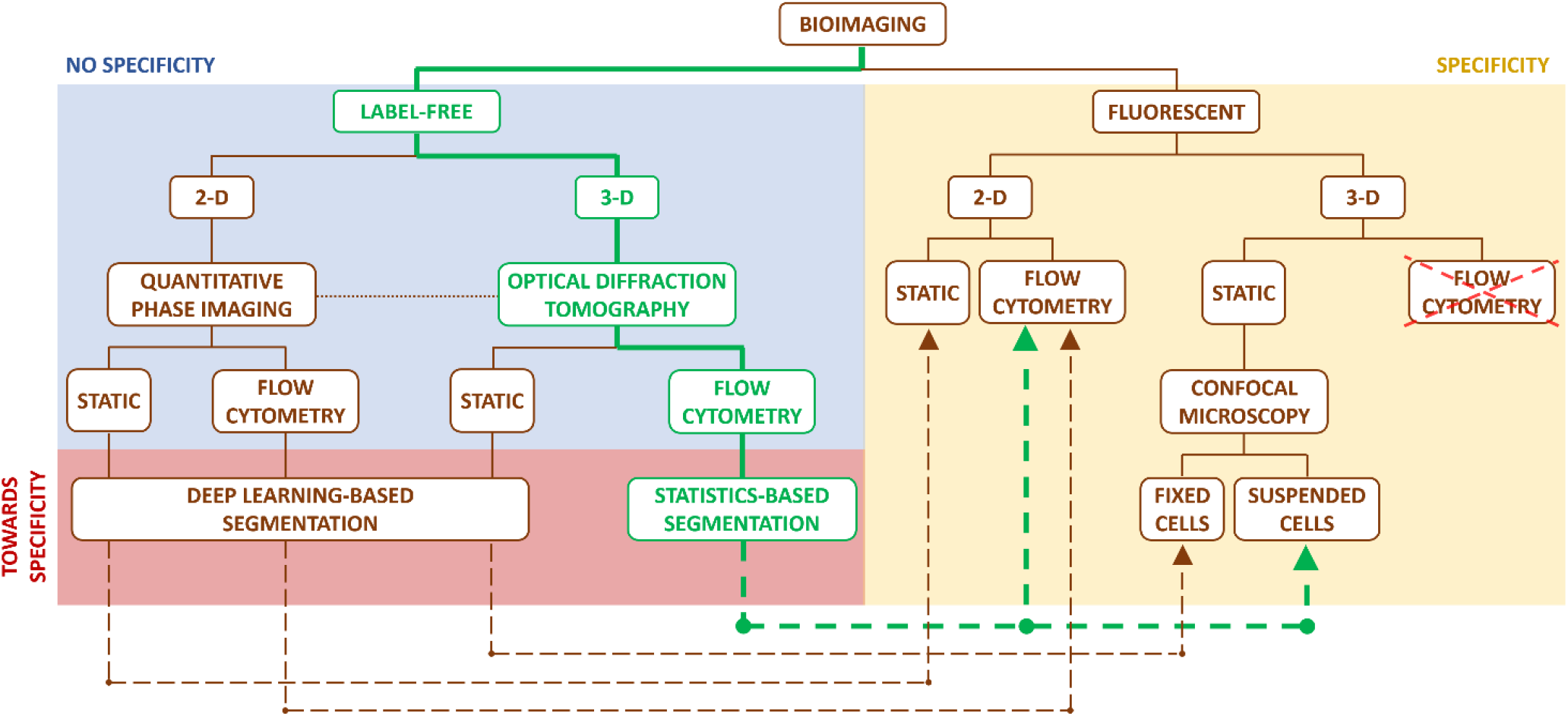
Comparison between label-free and fluorescent bioimaging in microscopy. Unlike the label-free bioimaging (blue box), the fluorescent bioimaging (yellow box) has sub-cellular specificity because nucleus is marked, but it is qualitative and limited by the staining itself. The methods in the red box allow to fill the specificity gap between the label-free and fluorescent techniques (dashed lines). In particular, a Generative Adversarial Network has been employed to virtually stain unlabelled tissues (PhaseStain)^22^ as well as single cells (PICS^25^ and HoloStain^26^) in QPMs, i.e. in a 2D imaging case. Moreover, digital staining through the application of deep neural networks has been successfully applied to multi-modal multi-photon microscopy in histopathology of tissues^23^. A neural network has also been used to translate autofluorescence images into images that are equivalent to the bright-field images of histologically stained versions of the same samples, thus achieving virtual histological staining^24^. In the 3D imaging case, nuclei of unlabelled and adhered cells have been identified using a deep convolutional neural network^27^ to introduce specificity in ODT reconstructions, thus making 3D label-free ODT equivalent to the well-established 3D confocal microscopy, but only for static analysis of fixed cells at rest on a surface. Instead, the specificity property of confocal microscopy has not been replicated yet on suspended cells in a label-free manner. The proposed technology (green pathway) fills a blank in the bioimaging realm because, in terms of specificity, the statistics-based segmentation (CSSI method) makes the holographic learning flow cyto-tomography consistent with both the 2D FM flow-cytometry and the 3D FM confocal microscopy of suspended cells. Moreover, it emulates a non-existent FM technique, i.e. the in-flow confocal microscopy. The dashed green lines indicate the fluorescence techniques that we used to validate the accuracy of the proposed method, and they also point to which conventional fluorescence techniques could be replaced by our technique.

In this paper, we propose a new 3D shape retrieval method, named here as Computational Segmentation based on Statistical Inference (CSSI), to identify the nucleus-like region in 3D ODT reconstructions in flow cytometry. To this aim, the holographic learning flow cyto-tomography system is implemented by combining tomographic flow cytometry by digital holography (DH)^28,29^ and Learning Tomography (LT) reconstruction algorithm^30^. Our method is completely different than the others followed so far^22-27^, since it avoids the learning step and exploits a robust *ad hoc* clustering algorithm, i.e., it recognizes statistical similarities among the nucleus voxels. Furthermore, to the best of our knowledge, for the first time we are posing the problem of delineating the stain-free nucleus in 3D within single suspended cells flowing in a microfluidic cytometer. Beyond the numerical assessment through virtual 3D cell phantoms, we show that the CSSI algorithm can fill the specificity gap with two-dimensional (2D) FM cyto-fluorimetry and with 3D FM confocal microscopy. In addition, CSSI allows for direct measurements at the stain-free nuclear level of intrinsic 3D parameters (morphology, RI, and their derivatives, like dry-mass) correlated to cell physiology and health state^11^, thus providing a whole label-free quantitative characterization exploitable for analyzing large numbers of flowing single-cells^31^.

## Results

### Description and assessment of the CSSI method

To reconstruct the 3D RI distribution at the single-cell level in-flow, the 3D holographic learning flow cyto-tomography system sketched in Fig. 2a has been employed, which is described in the Methods section along with the numerical processing summarized in Fig. 2b. Since the sub-cellular structures (i.e. organelles) cannot be detected within the tomogram by conventional RI-based thresholding methods, the CSSI algorithm is proposed to identify the nucleus-like region.

**Fig. 2.**
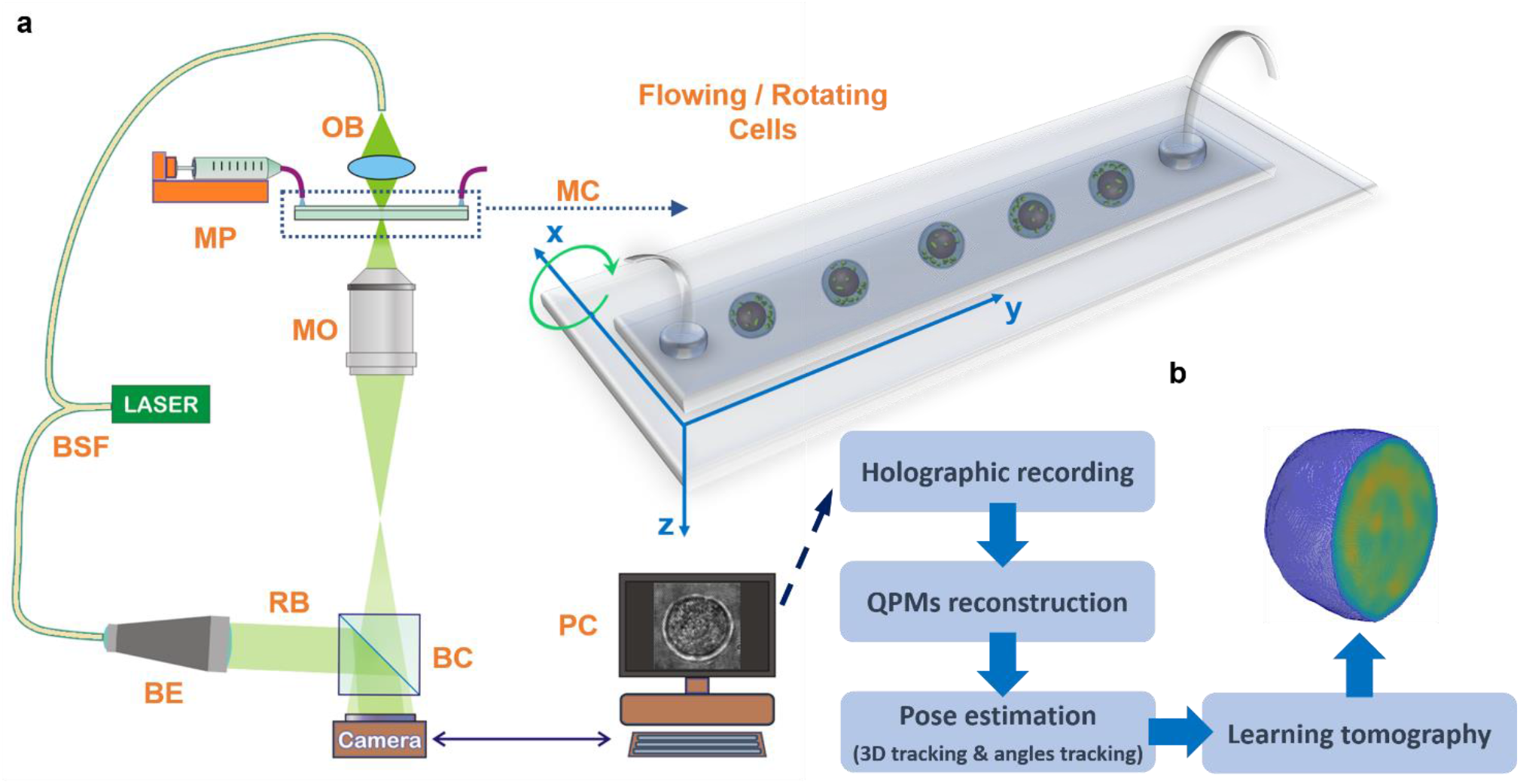
3D Holographic learning flow cyto-tomography technique. **a** Sketch of the in-flow ODT setup based on a DH microscope in off-axis configuration. BSF, Beam Splitter Fiber; MP, Microfluidic Pump; MC, Microfluidic Channel; OB, Object Beam; MO, Microscope Objective; BE, Beam Expander; RB, Reference Beam; BC, Beam Combiner; PC, Personal Computer. According to the reference system, cells flow along the y-axis, rotate around the x-axis, and are illuminated along the z-axis. **b** Block diagram of the holographic processing pipeline to reconstruct the stain-free 3D RI tomograms of flowing and rotating single-cells. Multiple digital holograms are recorded around the cell, from which the corresponding QPMs are numerically retrieved. The pose of each flowing cell is calculated by the 3D holographic tracking method and the rolling angles recovery approach. Finally, LT algorithm is exploited to enhance the initial tomographic reconstruction obtained by the inverse Radon transform.

To validate the CSSI method, we preliminarily tested and assessed it on a 3D numerical cell phantom simulation, modelled with the cell membrane, nucleus, cytoplasm, and mitochondria, as shown in Fig. 3a (see the Methods section). As reported by the histogram in Fig. 3b, a RI distribution has been assigned to each of the four sub-cellular structures. This 3D numerical cell phantom has been used to assess our proposed CSSI algorithm in the case of the nucleus segmentation. However, this is only the particular case of a more general technique which in principle can segment any kind of subcellular structure with a suitable spatial resolution, because it exploits the only hypothesis of knowing the location of a group of voxels belonging to the organelle to be segmented, considered as initial reference set. In fact, the CSSI method is based on the Wilcoxon–Mann–Whitney (WMW) test^32,33^, that is a statistical test we use to reject or not the hypothesis for which a test set has been drawn from the same distribution of a designed reference set. In particular, the steps depicted in the scheme in Fig. 3c are performed as follows

**Fig. 3.**
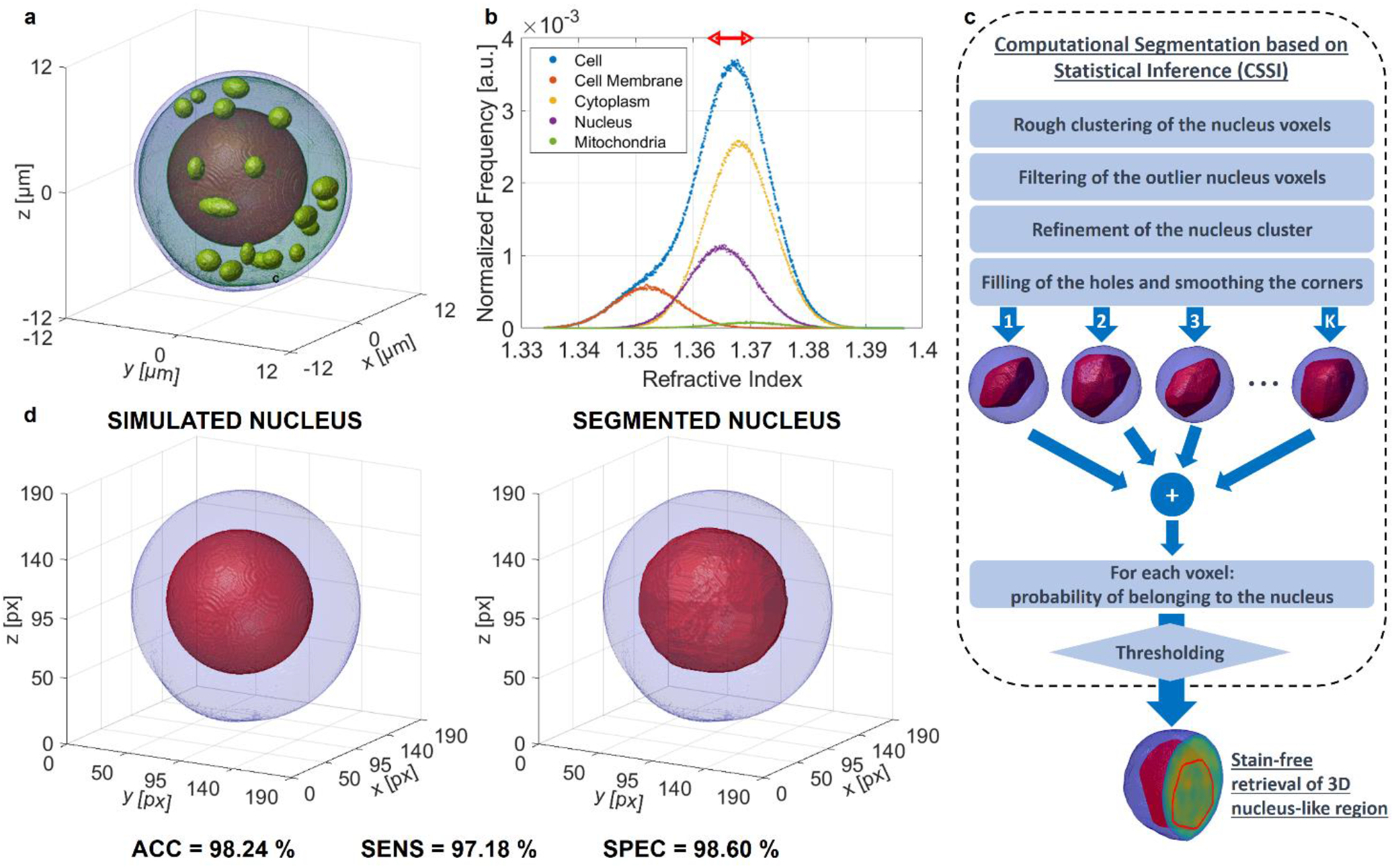
Numerical assessment of the CSSI algorithm applied to segment the 3D nucleus-like regions from a 3D numerical cell phantom (Supplementary Movie 1). **a** Isolevels representation of the 3D cell model, simulated with four sub-cellular components, i.e., cell membrane, cytoplasm, nucleus, and 18 mitochondria. **b** Histogram of the RI values assigned to each simulated sub-cellular structure in (a). The red arrow at the top highlights the RI values assigned to the transition region between the nucleus and cytoplasm. **c** Block diagram of the CSSI method to segment the nucleus-like region from a stain-free 3D RI tomogram. **d** Visual comparison between the simulated 3D nucleus and the 3D nucleus-like region segmented from the simulated RI tomogram in (a). The simulated nucleus and the segmented nucleus are marked in red within the blue cell shell. The clustering performances obtained in this simulation are reported below.

- Rough ε-clustering of the organelle voxels, exploiting the WMW test to infer the statistical similarity between different voxel clouds of side ε pixels (i.e. the test sets) and a certain voxel cloud (i.e. the initial reference set) which contains the voxels supposed belonging to the organelle of interest.
- Filtering of the outlier organelle voxels, because too far away from the centroid of the rough organelle cluster in terms of both geometric and statistical distances.
- Refinement of the filtered organelle cluster to improve its external shape by adding/removing smaller voxel clouds.
- Filling of the holes and smoothing of the corners of the refined organelle cluster by common morphological operators.

Notice that, whenever the WMW test is used, the reference set is randomly selected from the last estimation of the organelle cluster until that moment, to match its dimensionality with that of the test set, thus preserving the fairness of the statistical test. Due to this random selection, by repeating several times the described steps, at each iteration *j* = 1,2,…,*K* we can obtain a slightly different estimation of the organelle-like region. The output of each iteration is a binary 3D volume whose non-null values correspond to the voxels associated with the organelle. Therefore, the sum of all the *K* outputs provides a tomogram of occurrences, from which the probability that a voxel belongs to the organelle can be inferred through a normalization operation. Finally, the organelle-like region is identified by a suitable probability threshold. A detailed description of the CSSI algorithm is reported in the Supplementary Information. In this paper we focalize on the problem of the stain-free nucleus segmentation, therefore we associate the initial reference set to the central voxels of the cell. Indeed, for many kinds of suspended cells, the central voxels belong to the nucleus. This property occurs especially in the case of the cancer cells. In fact, it is confirmed by the 2D images of the neuroblastoma cells recorded through a FM cyto-fluorimeter (see Fig. S3), by the 3D morphological parameters reported in the literature for MCF-7 cells imaged through a confocal microscope^34^, and more generally by the increase of the nucleus-cytoplasm ratio demonstrated in cancer cells^35-39^.

On the left in Figure 3d, the sole simulated nucleus is reported in red within the blue cell shell, while on the right we show the nucleus segmented from the 3D numerical cell phantom through the CSSI algorithm. The visual comparison in Fig. 3d suggests that the proposed CSSI method allows segmenting a nucleus region very close to the original one, as also confirmed by the great quantitative performances reported below the tomograms, i.e., the accuracy (ACC), sensitivity (SENS), and specificity (SPEC) defined in the Methods section. Moreover, to numerically assess the proposed 3D CSSI algorithm, it has been applied to reconstruct the nuclei of 100 numerical cell phantoms simulated by randomly drawing their morphological and RI parameters from the distributions described in the Methods section and in Table S2. The overall average performances resulted to be very high, i.e. *ACC* = 96.46 %,*SENS* = 94.77 %, and *SPEC* = 97.08 %.

### Experimental consistency with 2D FM cyto-fluorimetry

The proposed CSSI method has been used to retrieve the 3D nucleus-like regions from five stain-free human neuroblastoma cancer cells (SK-N-SH cell line), reconstructed by 3D holographic learning flow cyto-tomography. The isolevels representation of an SK-N-SH cell is shown in Fig. 4a, highlighting in red the 3D segmented nucleus-like region within the blue cell shell. Moreover, its central slice is displayed in Fig. 4b, in which the segmented nucleus is marked by the red line, while in Fig. 4c we report in green the corresponding 3D RI histogram, separating in red and in blue the contributions of the 3D nucleus-like region and the 3D non-nucleus one, respectively. To experimentally assess the 3D segmentation technique, we digitally projected the segmented 3D ODT reconstruction back to 2D where the experimental 2D FM images are available for comparison. In particular, the segmented RI tomogram is digitally rotated from 0° to 150° with 30° angular step around *x*-, *y*-, and *z*-axes, and then its silhouettes along the *z*-, *x*-, and *y*-axes, respectively, are considered to create 2D ODT segmented projections, as sketched in Fig. 4a. According to the ray optics approximation, the phase measured by DH is directly proportional to the integral of the RI values along the direction perpendicular to the plane of the camera. In this way, 18 unlabelled QPMs were obtained. As shown in a re-projected QPM on the left in Fig. 4d, it is cumbersome to recognize a sub-cellular structuring since no label is employed. However, thanks to the proposed 3D CSSI algorithm, the region occupied by the nucleus can be also marked (red line) in the 2D ODT projection within the outer cell (blue line). We exploit this process to further assess the proposed segmentation algorithm, by making the 3D results obtained through the in-flow ODT technique comparable with a conventional 2D FM cyto-fluorimeter (i.e. ImageStreamX, see the Methods section). The latter has been used to record 1280 2D FM images of flowing SK-N-SH single cells, in which the nuclei have been stained through fluorescent dyes. On the right in Fig. 4d, the bright-field image of an SK-N-SH cell has been combined with the corresponding fluorescent image of the marked nucleus, therefore the false-color visualization makes the nucleus easily distinguishable (red line) with respect to the outer cell (blue line). ImagesStreamX can record a single random 2D image for each cell since it goes through the field of view (FOV) once. Instead, ODT allows the 3D tomographic reconstruction of a single cell. Through the reprojection process, we simulate the transition of the reconstructed cell within the ImageStreamX FOV at different 18 3D orientations with respect to the optical axis. In this way, we digitally replicate the ImageStreamX recording process and we also increase the dataset of 2D ODT images avoiding a high correlation between the reprojections of the same cell, thanks to the choise of a big angular step (i.e., 30°). Hence, in the 3D scatter plot in Fig. 4e, we quantitively compare some 2D morphological parameters representative of nucleus size, nucleus shape, and nucleus position, i.e. nucleus-cell area ratio (NCAR), nucleus aspect ratio (NAR), and normalized nucleus-cell centroid distance (NNCCD), respectively, measured from both 90 ODT images (red dots) and 1280 FM images (blue dots). In particular, we computed the NAR as the ratio between the minor axis and the major axis of the best-fitted ellipse to the nucleus surface, while the nucleus-cell centroid distance refers to 2D centroids and has been normalized to the radius of a circle having the same area of the cell, thus obtaining NNCCD. The 3D scatter plot highlights the great matching between ODT and FM 2D nuclear features since the ODT red dots are completely contained within the FM blue cloud. In addition, by using the WMW test between ODT and FM measurements about NCAR, NAR, and NNCCD, we also obtained high p-values, i.e., 0.980, 0.917, and 0.841, respectively, according to which it is not rejected with high confidence level the hypothesis that ODT and FM 2D nuclear features have been drawn from the same distributions. This quantitative comparison is summarized in Table S3. Moreover, to better visualize the 3D scatter plot in Fig. 4e, we split it into three different 2D scatter plots, shown in Figs. 4f-h.

**Fig. 4.**
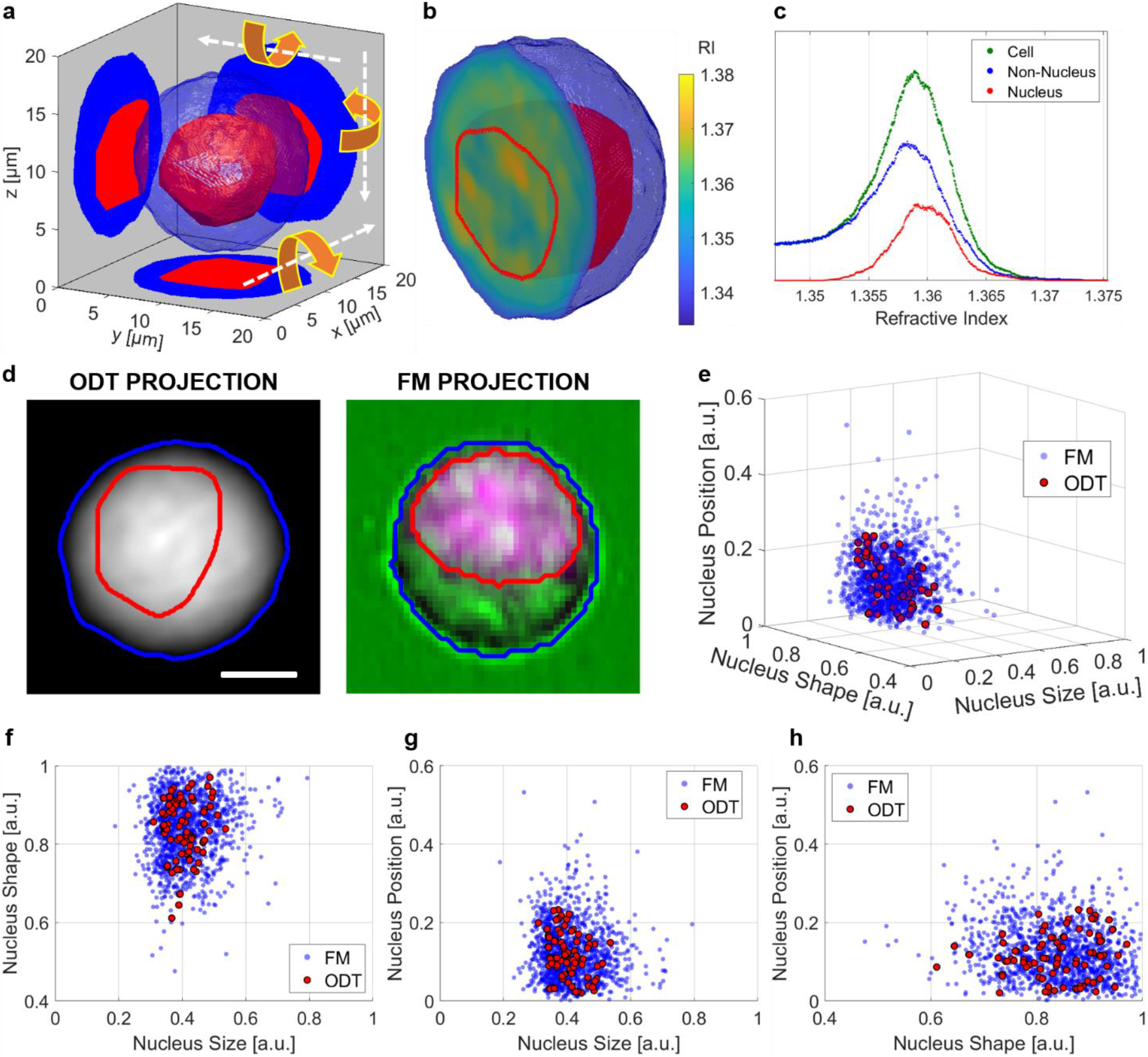
Experimental assessment of the CSSI algorithm applied to segment the 3D nucleus-like regions from unlabelled in-flow ODT reconstructions of five SK-N-SH cells, by comparison with the morphological parameters measured through a 2D FM cyto-fluorimeter (Supplementary Movie 2). **a** 3D segmented nucleus (red) within the 3D cell shell (blue) of an SK-N-SH cell reconstructed by ODT. The segmented tomogram is rotated around the *x*-, *y*-, and *z*-axes (orange arrows) and then reprojected along the *z*-, *x*-, and *y*-axes (white arrows), thus obtaining 2D ODT segmented projections in *xy*-, *yz*-, and *xz*-planes, respectively. **b** Central slice of the isolevels representation in (a), with nucleus marked by the red line. **c** RI histogram of the SK-N-SH cell in (a,b) reconstructed by 3D in-flow ODT (green), along with the RI distributions of its 3D nucleus-like region (red) and non-nucleus one (blue) segmented by CSSI algorithm. **d** 2D segmented projection with nucleus (red line) and non-nucleus (blue line) regions, obtained (on the left) by reprojecting 3D unlabelled ODT RI reconstruction in (a,b) and (on the right) by recording 2D labelled FM images. The scale bar is 5 μm. **e** 3D scatter plot of nucleus size vs nucleus shape vs nucleus position measured in 1280 FM (blue dots) and 90 ODT (red dots) 2D projections. Nucleus size is NCAR, nucleus shape is NAR, and nucleus position is NNCCD. **f-h** 2D scatter plots of nucleus size vs nucleus shape, nucleus size vs nucleus position, and nucleus shape vs nucleus position, respectively, containing the same points in (e).

### Experimental consistency with 3D FM confocal microscope

For the second experimental assessment, three stain-free human breast cancer cells (MCF-7 cells) have been reconstructed by 3D holographic learning flow cyto-tomography and then segmented by the CSSI method, as shown in the example in Figs. 5a,b. In particular, the nucleus shell is marked in red within the blue cell shell in the isolevels representation of Fig. 5a, which segmented central slice is displayed in Fig. 5b. Moreover, in Figure 5c, we display in green its 3D RI histogram, also separating the RI distribution of the 3D nucleus-like region (red) and the 3D non-nucleus one (blue). In this case, the experimental assessment is based on a quantitative comparison with the 3D morphological parameters measured in ref. 34, in which a confocal microscope has been employed to find differences between viable and apoptotic MCF-7 cells through 3D morphological features extraction. In this study, 206 suspended cells were stained with three fluorescent dyes to measure average values and standard deviations of 3D morphological parameters about the overall cell and its nucleus and mitochondria. A synthetic description of 3D nucleus size, shape, and position is given by nucleus-cell volume ratio (NCVR), nucleus surface-volume ratio (NSVR), and normalized nucleus-cell centroid distance (NNCCD), respectively. In particular, in this case, the nucleus-cell centroid distance refers to 3D centroids and has been normalized with respect to the radius of a sphere having the same cell volume, thus obtaining NNCCD. Moreover, it is worth underlining that NCVR and NSVR are direct measurements reported in ref. 34, while NNCCD is an indirect measurement since it has been computed by using the direct ones in ref. 34. In the 2D scatter plots in Figs. 5d-f regarding nucleus size, shape, and position, the three ODT measurements (red dots) are reported along with three blue rectangles, which are the intervals *µ* ±1σ, *µ* ±2σ, and *µ* ±3σ, with *µ* the average value and σ the standard deviation of the same parameters measured by FM confocal microscope. These scatter plots highlight a very good agreement between the 3D nucleus identified in labelled static MCF-7 cells by confocal microscope and the 3D nucleus segmented in unlabelled flowing MCF-7 cells by the proposed CSSI algorithm. In fact, all the ODT values are located in the 1σ-interval around the FM average values, except for shape measurement, that is anyway located in the 2σ-interval around the FM average value (Figs. 5d,f). The values shown in Figs. 5d-f are summarized in Table S4.

**Fig. 5.**
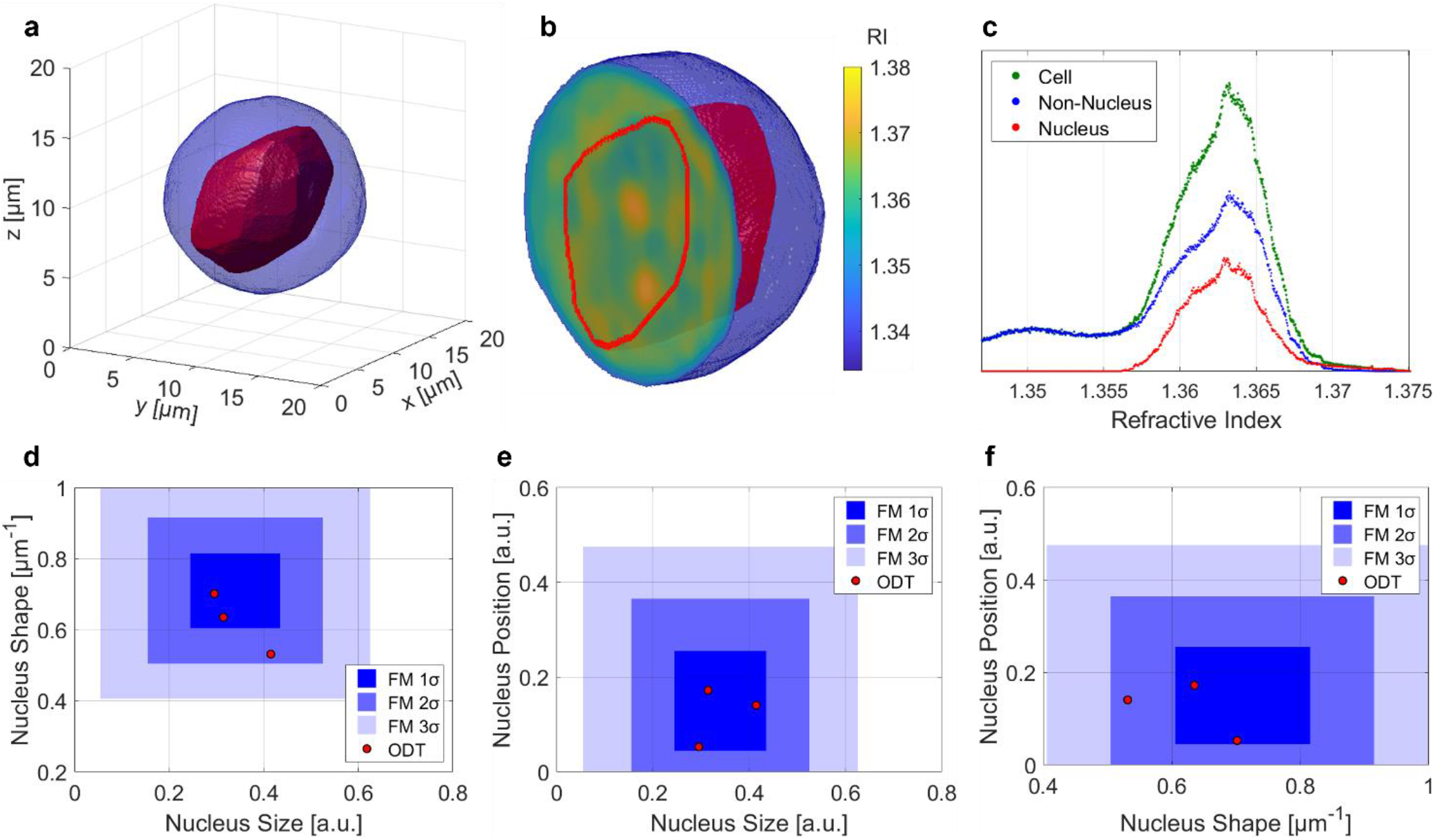
Experimental assessment of the CSSI algorithm applied to segment the 3D nucleus-like regions from unlabelled in-flow ODT reconstructions of three MCF-7 cells, by comparison with the morphological parameters measured through a 3D FM confocal microscope (Supplementary Movie 2). **a** 3D segmented nucleus-like region (red) within unlabelled 3D cell shell (blue) reconstructed through in-flow ODT. **b** Central slice of the isolevels representation in (a), with nucleus marked by the red line. **c** RI histogram of the MCF-7 cell in (a,b) reconstructed by 3D in-flow ODT (green), along with the RI distributions of its 3D nucleus-like region (red) and non-nucleus one (blue) segmented by CSSI algorithm. **d-f** Scatter plots of nucleus size vs nucleus shape, nucleus size vs nucleus position, and nucleus shape vs nucleus position, respectively, measured in three segmented ODT MCF-7 nuclei (red dots) along with the corresponding FM intervals (blue rectangles) around the average values, with half-width 1σ, 2σ, and 3σ (σ is the standard deviation of the measurements). Nucleus size is NCVR, nucleus shape is NSVR, and nucleus position is NNCCD.

## Methods

### Sample preparation

The human breast cancer cells (MCF-7 cell line) and the human neuroblastoma cells (SK-N-SH cell line) were selected for tomographic experiments. The MCF-7 and the SK-N-SH cells were cultured in RPMI 1640 and in Minimum Essential Medium Eagle (MEM) from Sigma Aldrich, respectively. Both cell culture media were supplemented with 10% fetal bovine serum, 2 mM L-glutamine, 100 μg ml^-1^ streptomycin, and 100 U ml^-1^ penicillin. Then, cells were collected from the Petri dish by incubation for 5 min with a 0.05% trypsin–EDTA solution (Sigma, St. Louis, MO). Finally, the MCF-7 and the SK-N-SH cells were centrifuged for 5 min at 1500 rpm and 1200 rpm, respectively, resuspended in complete medium, and injected into the microfluidic channel for the holographic flow cyto-tomography experiment. The SK-N-SH cells were also analysed through a FM cyto-fluorimeter. To this aim, 7 million SK-N-SH cells were resuspended in Phosphate Buffered Saline (PBS), 1X (Sigma) and exposed to 25 µM DRAQ5 Fluorescent Probe for 5 min at room temperature under agitation (#62254, Thermo Scientific™).

### Holographic flow cyto-tomography setup

In a QPM, phase-contrast is due to the optical path length difference between the unlabelled biological specimen and its background because of the combination of its thickness and its RI^2^. These two quantities can be decoupled by recording multiple 2D QPMs at different viewing angles around the sample, thus performing the 3D ODT. However, unlike conventional ODT methods, in our setup the DH microscope acquires multiple digital holograms of flowing and rotating cells within a microfluidic channel, exploiting the hydrodynamic forces produced by a laminar flow^28,29^. In fact, the light beam generated by the laser (Laser Quantum – Torus, emitting at a wavelength *λ* = 532) is coupled into an optical fibre, which splits it into an object beam and a reference beam in order to constitute a Mach-Zehnder interferometer in off-axis configuration. The object beam exits from the fibre and is collimated to probe the biological sample that flows at 7 nL/s along a commercial microfluidic channel with cross section 200 μm × 200 μm (Microfluidic Chip-Shop). The flux velocity is controlled by a pumping system (CETONI – neMESYS) that ensures temporal stability of the parabolic velocity profile into the microchannel. The wavefield passing throughout the sample is collected by the Microscope Objective (Zeiss 40× – oil immersion – 1.3 numerical aperture) and directed to the 2048 × 2048 CMOS camera (USB 3.0 U-eye, from IDS) by means of a Beam-Splitter that allows the interference with the reference beam. The interference patterns of the single cells rotating into a 170 μm × 170 μm FOV are recorded at 35 fps.

According to the reference system sketched in Fig. 2a, cells flow along the y-axis and continuously rotate around the x-axis thanks to the microfluidic properties^45^, while their holograms are recorded along the z-axis. Then, as summarized in Fig. 2b, numerical operations are performed to reconstruct the stain-free 3D RI tomograms of the recorded flowing single-cells. The off-axis configuration allows demodulating each hologram of the recorded sequence by filtering the real diffraction order^46^. Then, the 3D positions of the flowing cells within the microfluidic channel are computed through a holographic tracking algorithm^47^. Each demodulated hologram is numerically propagated along the z-axis through the Angular Spectrum formula^46^ in order to minimize a contrast-based metric, i.e. the Tamura Coefficient^47^ (TC), to recover the z-position of the cell and refocus it. The QPMs are then obtained by implementing the phase unwrapping algorithm^46^ on the corresponding refocused complex wavefronts. In each QPM, the weighted centroids provide the *xy*-positions of the cell, which are used to center it in its cropped region of interest (ROI), as shown in Fig. 4d, thus avoiding motion artefacts in the final 3D tomogram. Moreover, the microfluidic properties and the high frame rate allow to linearize the relationship between the angular and the translational speeds^45^. Therefore, the *K* unknown rolling angles *ϑ*_*k*_ are estimated by using the computed *y*-positions as follows

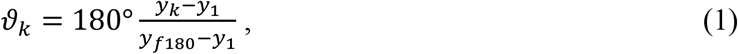

where *k* = 1, …, *K* is the frame index. The *f*_180_ value is the index of the frame at which the cell has rotated of 180° with respect to the first frame of the sequence. It is computed by minimizing the Tamura Similarity Index (TSI), that is a phase image similarity metric based on the evaluation of the local contrast calculated on all the QPMs of the rolling cell though the TC. The tomographic reconstruction is firstly obtained by the inverse Radon transform, and it is then enhanced though the LT algorithm.

### Learning Tomography algorithm

LT is an iterative reconstruction algorithm based on a nonlinear forward mode, beam propagation method (BPM), to capture high orders of scattering^30^. Using the BPM, we propagate an incident light illumination on an initial guess acquired by the inverse Radon transform and compare the resulting field with the experimentally recorded field. The error between the two fields is backpropagated to calculate the gradient^48^. At each iteration of LT, the gradient calculation is repeated for 8 randomly selected rotation angles, and the corresponding gradients are rotated and summed to update the current solution. As an intermediate step, the total variation regularization was employed. The total iteration number is 200 with a step size of 0.00025 and a regularization parameter of 0.005. In order to run LT, we need two electric fields, incident and total electric fields. The amplitude of the incident field was estimated from the amplitude of the total electric field by low-pass filtering in the Fourier domain with a circular aperture whose radius is 0.176 *k*_0_, where 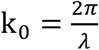 given a wavelength *λ* = 532 nm in a vacuum. In other words, we assume that the high-frequency information in the amplitude of the total electric field was only attributed to the light interference caused by a sample when illuminated by an illumination with slowly varying amplitude.

### FM cyto-fluorimeter

In order to record thousands of 2D FM images, a commercial multispectral flow cyto-fluorimeter has been employed, i.e., Amnis ImageStreamX®. Cells are hydrodynamically focused within a micro-channel, and then they are probed both by a transversal brightfield light source and by orthogonal lasers. The fluorescence emissions and the light scattered and transmitted from the cells are collected by an objective lens. After passing through a spectral decomposition element, the collected light is divided into multiple beams at different angles according to their spectral bands. The separated light beams propagate up to 6 different physical locations of one of the two CCD cameras (256 rows of pixels), which operates in time operation. Therefore, the image of each single flowing cell is decomposed into 6 separate sub-images on each of the two CCD cameras, based on their spectral band, thus allowing the simultaneous acquisition of up to 12 images of the same cell, including brightfield, scatter, and multiple fluorescent images. Hence, Amnis ImageStreamX® combines the single-cell analysis of the standard FM microscopy with the statistical significance due to large number of samples provided by standard flow-cytometry.

ImageStreamX Mark II Flow Cytometer (Luminex Corporation) was used to acquire 1280 single cells images at 60x magnification. For each single cell, we recorded two simultaneous images, i.e., a brightfield image of the flowing cell and its corresponding FM image. To segment nucleus, we analysed the single cells using the IDEAS software (version 6.2.64.0), which combines fluorescence intensity (DRAQ5) and morphometric measures (darkfield), generating a global threshold of the FM signal corresponding to the nucleus size. The automatic calibration provided by the instrument, corresponding to 0.33 μm/pixel, was employed to measure 2D morphological parameters related to the cell and the nucleus, in order to realize a quantitative comparison with the tomographic reconstructions. Three of the recorded brightfield and fluorescent images are shown at the top and at the bottom of Fig. S3, respectively, with overlapped in red the contour of the segmented nuclei.

### 3D numerical cell phantom

In ref. 34, a confocal microscope has been employed to find differences between viable and apoptotic MCF-7 cells through 3D morphological features extraction. In particular, 206 cells were stained with three fluorescent dyes in order to measure the average value and standard deviation of 3D morphological parameters about the overall cell and its nucleus and mitochondria. We exploit these measurements to simulate a 3D numerical cell phantom, by setting 1 px = 0.12 μm. It is made of four sub-cellular structures, i.e., cell membrane, cytoplasm, nucleus, and mitochondria. We shape cell, nucleus, and mitochondria as ellipsoids, then we make irregular the cell external surface, and finally we obtain cytoplasm through a morphological erosion of the cell shape. Moreover, in each simulation, the number of mitochondria is drawn from the uniform distribution *U*_1_ {*a*_1_, *b*_1_ }. A 3D numerical cell phantom is displayed in Fig. 3a, in which 18 mitochondria have been simulated. To each simulated 3D sub-cellular component, we assign a RI distribution, as shown by the RI histogram in Fig. 3b. Measuring accurate RI values at sub-cellular level is still a deeply debated topic^49,50^. Hence, we cannot replicate realistic RIs since they are not yet well known, therefore we simulate the unfavourable case for our testing purpose segmenting the nucleus from cytoplasm, i.e., we model overlapped subcellular distributions of the RI values. In particular, for each cell membrane voxel, we draw its RI from distribution *N*_1_ (*µ*_1_,*σ*^2^). Instead, without knowing if the nucleus RIs are greater than the cytoplasm ones or vice versa, in each simulation we randomly assign cytoplasm and nucleus to distributions *N*_2_ (*µ*_2_,*σ*^2^) or *N*_3_ (*µ*_3_,*σ*^2^). It is worth remarking that, to strengthen the numerical assessment, we increase the randomness of the RI assignments among the different simulations, because each voxel belonging to cell membrane, nucleus, and cytoplasm is drawn from gaussian distributions *N*_1_, *N*_2_, and *N*_3_ (or *N*_1_, *N*_3_, and *N*_2_), respectively, which average values *µ*_1_, *µ*_2_, and *µ*_3_ are in turn drawn from other gaussian distributions for each voxel extraction, i.e. 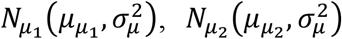 and 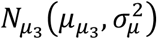, respectively. Instead, as regards mitochondria, each of them has a RI gaussian distribution *N*_4_ (*µ*_4_,*σ*^2^) which average value *µ*_4_ is drawn from the gaussian distribution 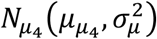 for each mitochondrion and not for each voxel. Moreover, we create a RI transition zone straddling the nucleus to cytoplasm, thus avoiding any discontinuity that could somehow facilitate the segmentation. In particular, after drawing all nucleus and cytoplasm values, RIs that are in the middle of their average values are assigned to the voxels of the transition zone, as highlighted by the red arrow at the top of Fig. 3b. This transition zone is obtained through morphological erosion and dilation of the nucleus ellipsoid, by using a spherical structuring element, which radius is drawn from the uniform distribution *U*_1_ {*a*_2_, *b*_2_ } px for each simulation, thus resulting in an internal nucleus volume that is about 85-95 % of the total nucleus volume. In the example in Figs. 3a,b, a 3 px radius has been selected. All the described parameters are reported in Table S2.

In order to quantify the performances of the CSSI algorithm, we compute the accuracy, sensitivity, and specificity respectively as follows

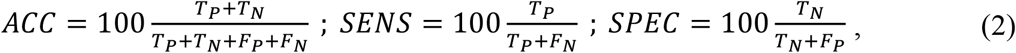

where TP (True Positive) is the number of voxels that are correctly classified as nucleus, TN (True Negative) is the number of voxels that are correctly classified as non-nucleus, FP (False Positive) is the number of voxels that are wrongly classified as nucleus, and FN (False Negative) is the number of voxels that are wrongly classified as non-nucleus. Therefore, accuracy is the percentage of voxels correctly classified, sensitivity is the percentage of voxels correctly classified nucleus with respect to the number of nucleus voxels, and specificity is the percentage of voxels correctly classified non-nucleus with respect to the number of non-nucleus voxels.

## Discussion

In this paper, we introduce and discuss a wholly new strategy for bridging the gap between FM and ODT in terms of subcellular specificity. In particular, we demonstrate for the first time the capability in identifying the cell nucleus from 3D phase-contrast tomograms in unstained cells analysed in a flow cytometry modality. Exploiting the learning-based tomographic reconstruction algorithm, we developed the CSSI method to identify a stain-free organelle region. Direct experimental validation of the proposed CSSI method is very difficult in flow-cytometry and too prone to errors that could alter the truthfulness of the experiment, as the cells should be stained and recorded simultaneously by both the holographic and the fluorescent channels. The numerical simulations provided great performances in accomplishing the task of nuclei extraction and measurement by our new proposed approach. In addition, we performed the CSSI experimental assessment by retrieving the stain-free nuclei for two distinct cancer cell lines starting from the 3D holographic learning flow cyto-tomograms, and we discussed the consistency with the classical FM microscopic methods (see green dashed lines in Fig. 1).

To provide a general overview of the differences, advantages, and limitations of the main works that aim at introducing nucleus specificity, in Table 1 we compare the most significant efforts towards this task. Each cell of Table 1 is filled with flags or crosses depending on whether the corresponding method possesses or lacks a certain attribute, each of them being highly pursued in the bioimaging field. PhaseStain^22^, PICS^25^, and HoloStain^26^ are virtual staining-based methods which use AI (Generative Adversarial Networks) to emulate fluorescence in a stain-free manner, which is certainly one of the most interesting approaches shown lately to overcome the main limitation of QPI towards the organelle specificity. However, they intrinsically rely on 2D fluorescence for training an AI, and the results strictly depend on the training dataset. In the 3D case, the most promising approach to add specificity to ODT is based on a Convolutional Neural Network^27^, which so far has been validated by identifying cells’ nuclei in label-free modality for adhering samples. Again, the approach relies on AI to segment the nucleus. In this sense, we believe the CSSI method is very promising to promote label-free ODT with nucleus specificity since it addresses all the required attributes shown in Table 1. In particular, the proposed CSSI algorithm allows in-flow ODT to reach the same results of 2D FM cyto-fluorimeter, but without using dyes and preserving its high-throughput property. Furthermore, the ODT reprojections are much more informative than the FM images (Fig. 4d). Indeed, the phase values contain a quantitative measurement about both the 3D sub-cellular morphology and RI distribution, which can be associated to the cell biology, instead of the 2D FM images, from which the sole 2D morphological parameters can be inferred. Similarly, besides the 3D morphological analysis of the confocal microscopy, in our technology a complete 3D label-free quantitative characterization of the RI-based fingerprint at the sub-cellular single-cell level is possible, as reported in the histograms in Fig. 4c and Fig. 5c. As demonstrated in the present paper, the CSSI approach works properly with any type of cells since it exploits an *ad hoc* clustering algorithm which segments the nucleus-like region thanks to the statistical similarities of its voxels. CSSI strength lies in completely surpassing the limitation of AI-based techniques that exploit neural networks previously trained through FM images and can only supply results about the specific situations they have been exposed to during the learning process. The second fundamental aspect is that the CSSI method retrieves the 3D nuclear specificity in stain-free suspended single-cells in flow cytometry, thus providing quantitative measurements at the sub-cellular level with statistical significance on a large number of cells by potentially exploiting the high-throughput property (not possible through a confocal microscope). Finally, the same CSSI algorithm can be also applied for the segmentation of any kind of stain-free organelle with a suitable spatial resolution by just changing the initial reference set. Future experiments will be dedicated to this aim. In conclusion, we believe that, thanks to the above-mentioned properties, the CSSI algorithm combined with the holographic learning flow cyto-tomography could open a new route for label-free microscopy as biomedical tool. By starting from this conceptually breaking strategy, such a tool could revolutionize the cancer diagnosis^40-42^ (e.g., through the liquid biopsy paradigm^43^) or more in general affirm ODT as a viable method for intracellular quantitative characterization at the single-cell level, which can be further exploited for therapeutic purposes in personalized medicine^44^.

**Table 1.** Properties of the methods for the nucleus identification.

| **Table 1.** Properties of the methods for the nucleus identification. |  |  |  |  |
| --- | --- | --- | --- | --- |
|  | **Label-free** | **3D** | **Flow Cytometry** | **Specificity** |
| **QPI** | ✓ | ✗ | ✓ | ✗ |
| **ODT** | ✓ | ✓ | ✓ | ✗ |
| **FM Cytometry** | ✗ | ✗ | ✓ | ✓ |
| **FM Confocal Microscopy** | ✗ | ✓ | ✗ | ✓ |
| **PhaseStain<sup>22</sup> – PICS<sup>25</sup>** | ✓ | ✗ | ✗ | ✓ |
| **HoloStain<sup>26</sup>** | ✓ | ✗ | ✓ | ✓ |
| **ODT + Deep Learning** | ✓ | ✓ | ✗ | ✓ |
| **ODT + CSSI** | ✓ | ✓ | ✓ | ✓ |

## Acknowledgements

This work was funded at CNR-ISASI by project PRIN 2017, Morphological Biomarkers for early diagnosis in Oncology (MORFEO) Prot. 2017N7R2CJ.

This work was funded at EPFL by Swiss National Science Foundation (514481).

J.L. is partially funded by the Innosuisse project (34247.1 IP-ENG).

## Author contributions

L.M. and F.M. designed and conducted the holographic experiments; M.M., F.C. and A.M. took care of cell culture; D.Pi., P.M., J.L., V.B. designed and performed the numerical analysis. M.C. and A.I. contributed to conceive the experiments from biomedical point of view and validated the results of the experiments; F.V. and F.C. conducted the cytofluorimeter measurements and provided data; D.Ps. and P.F. coordinated and guided the research.

D.Pi., P.M., L.M., V.B., J.L., P.F., D.Ps. prepared the manuscript.

All authors supported data analysis and interpretation.

All authors reviewed the manuscript.

## Supplementary information

**Supplementary Movie 1 –** Steps of the CSSI algorithm for the nucleus segmentation in the 3D numerical cell phantom (MP4).

**Supplementary Movie 2 -** Holographic learning flow cyto-tomography and 3D nucleus segmentation by the CSSI algorithm in an MCF-7 cell and in an SK-N-SH cell (MP4).

## References

1. Lichtman, J. & Conchello, J. A. Fluorescence Microscopy. Nat. Methods 2, 910–919 (2005).

2. Park, Y., Depeursinge, C. & Popescu, G. Quantitative phase imaging in biomedicine. Nat. Photonics 12, 578–589 (2018).

3. Merola, F. et al. Diagnostic tools for lab-on-chip applications based on coherent imaging microscopy. Proc. IEEE 103(2), 192–204 (2015).

4. Cotte, Y. et al. Marker-free phase nanoscopy. Nat. Photonics 7, 113–117 (2013).

5. Yoffe, G. D., Mirsky, S. K., Barnea, I. & Shaked, N. T. High-resolution 4-D acquisition of freely swimming human sperm cells without staining. Sci. Adv. 6(15), eaay7619 (2020).

6. Kemper, B. et al. Label-free quantitative cell division monitoring of endothelial cells by digital holographic microscopy. J. Biomed. Opt. 15(3), 036009 (2010).

7. Zhang, Y. et al. Motility-based label-free detection of parasites in bodily fluids using holographic speckle analysis and deep learning. Light Sci. Appl. 7(1), 1–18 (2018).

8. Jo, Y. et al. Quantitative Phase Imaging and Artificial Intelligence: A Review. IEEE J Sel Top Quantum Electron 25(1), 1–14 (2019).

9. Zhang, J. K., He, Y. R., Sobh, N. & Popescu, G. Label-free colorectal cancer screening using deep learning and spatial light interference microscopy (SLIM). APL Photonics 5(4), 040805 (2020).

10. Kim, G., Jo, Y., Cho, H., Min, H. S. & Park, Y. K. Learning-based screening of hematologic disorders using quantitative phase imaging of individual red blood cells. Biosens Bioelectron 123, 69–76 (2019).

11. Zangle, T. A. & Teitell, M. A. Live-cell mass profiling: an emerging approach in quantitative biophysics. Nature Methods 11(12), 1221–1228 (2014).

12. Jin, D., Zhou, R., Yaqoob, Z. & So, P. Tomographic phase microscopy: Principles and applications in bioimaging. J. Opt. Soc. Am. B 34(5), B64–B77 (2017).

13. Kim, Y. et al. Profiling individual human red blood cells using common-path diffraction optical tomography. Sci. Rep. 4, 6659 (2014).

14. Kuś, A., Dudek, M., Kemper, B., Kujawińska, M. & Vollmer, A. Tomographic phase microscopy of living three-dimensional cell cultures. J. Biomed. Opt. 19, 046009 (2014).

15. Sung, Y., Choi, W., Lue, N., Dasari, R. R. & Yaqoob, Z. Stain-Free Quantification of Chromosomes in Live Cells Using Regularized Tomographic Phase Microscopy. PLoS ONE 7(11), e49502 (2012).

16. Yoon, J. et al. Label-free characterization of white blood cells by measuring 3D refractive index maps. Biomed. Opt. Express 6, 3865–3875 (2015).

17. Kim, K. et al. Three-dimensional label-free imaging and quantification of lipid droplets in live hepatocytes. Sci. Rep. 6, 36815 (2016).

18. Kim, K. et al. Optical diffraction tomography techniques for the study of cell pathophysiology. J. Biomed. Photonics Eng. 2, 020201 (2016).

19. Pirone, D. et al. Three-Dimensional Quantitative Intracellular Visualization of Graphene Oxide Nanoparticles by Tomographic Flow Cytometry. Nano Lett. 21, 5958–5966 (2021).

20. Wang, Z. et al. Dehydration of plant cells shoves nuclei rotation allowing for 3D phase-contrast tomography. Light.: Sci. Appl. 10, 187 (2021).

21. Kim, D. et al. Refractive index as an intrinsic imaging contrast for 3-D label-free live cell imaging. BioRxiv (2017).

22. Rivenson, Y. et al. PhaseStain: the digital staining of label-free quantitative phase microscopy images using deep learning. Light.: Sci. Appl. 8, 23 (2019).

23. Borhani, N., Bower, A. J., Boppart, S. A., & Psaltis, D. Digital staining through the application of deep neural networks to multi-modal multi-photon microscopy. Biomed. Opt. Express 10, 1339–1350 (2019).

24. Rivenson, Y. et al. Deep learning-based virtual histology staining using auto-fluorescence of label-free tissue. 1803.11293 (2018)

25. Kandel, M. E. et al. Phase imaging with computational specificity (PICS) for measuring dry mass changes in sub-cellular compartments. Nat. Commun. 11, 6256 (2020).

26. Nygate, Y. N. et al. Holographic virtual staining of individual biological cells. Proc. Natl. Acad. Sci. USA 117, 9223–9231 (2020).

27. Lee, J. et al. Deep-Learning-Based Label-Free Segmentation of Cell Nuclei in Time-Lapse Refractive Index Tomograms. IEEE Access 7, 83449–83460 (2019).

28. Merola, F. et al. Tomographic flow cytometry by digital holography. Light Sci. Appl. 6, e16241 (2017).

29. Villone, M. M. et al. Full-angle tomographic phase microscopy of flowing quasi-spherical cells. Lab Chip 18(1), 126–131 (2018).

30. Lim, J., Goy, A., Shoreh, M. H., Unser, M. & Psaltis, D. Learning tomography assessed using Mie theory. Phys. Rev. Appl. 9(3), 034027 (2018).

31. Liu, Y. et al. Cell Refractive Index for Cell Biology and Disease Diagnosis: Past, Present and Future. Lab Chip 16, 634–644 (2016).

32. Mann, H. B. & Whitney, D. R. On a test of whether one of 2 random variables is stochastically larger than the other. Annals of Mathematical Statistics 18, 50‐60 (1947).

33. Wilcoxon, F. Individual comparisons by ranking methods. Biometrics Bulletin 1, 80‐83 (1945).

34. Wen, Y. et al. Quantitative analysis and comparison of 3D morphology between viable and apoptotic MCF-7 breast cancer cells and characterization of nuclear fragmentation. PLoS ONE 12(9), e0184726 (2017).

35. McIntire, P. J. et al. Digital image analysis supports a nuclear-tocytoplasmic ratio cutoff value below 0.7 for positive for high-grade urothelial carcinoma and suspicious for high-grade urothelial carcinoma in urine cytology specimens. Cancer Cytopathol. 127(2), 120–124 (2019).

36. Liu, J. et al. Machine learning of diffraction image patterns for accurate classification of cells modeled with different nuclear sizes. Journal of Biophotonics 13(9), e202000036 (2020).

37. Moore, M. J., Sebastian, J. A. & Kolios, M. C. Determination of cell nucleus-to-cytoplasmic ratio using imaging flow cytometry and a combined ultrasound and photoacoustic technique: a comparison study. J. Biomed. Opt. 24, 106502 (2019).

38. Zink, D., Fischer, A. H. & Nickerson, J. A. Nuclear structure in cancer cells. Nat. Rev. Cancer 4, 677–687 (2004).

39. Takaki, T. et al. Actomyosin drives cancer cell nuclear dysmorphia and threatens genome stability”, Nat. Commun. 8, 16013 (2017).

40. Backman, V. et al. Detection of preinvasive cancer cells. Nature 406, 35–36 (2000).

41. Uttam, S. et al. Early Prediction of Cancer Progression by Depth-Resolved Nanoscale Mapping of Nuclear Architecture from Unstained Tissue Specimens. Cancer Res 75, 4718–4727 (2015).

42. Wang, P., Bista, R., Bhargava, R., Brand, R. E. & Liu, Y. Spatial-domain low-coherence quantitative phase microscopy for cancer diagnosis. Opt Lett 35(17), 2840–2842 (2010).

43. Miccio, L. et al. Perspectives on liquid biopsy for label free detection of “circulating tumor cells” through intelligent lab on chips. View 1(3), 20200034 (2020).

44. Sung, W. et al. Computational modeling and clonogenic assay for radioenhancement of gold nanoparticles using 3D live cell images. Radiat. Res. 190(5), 558–564 (2018).

45. Pirone, D. et al. Rolling angles recovery of flowing cells in holographic tomography exploiting the phase similarity. Appl Opt 60(4), A277–A284 (2021).

46. Kim, M. K. Principles and techniques of digital holographic microscopy. SPIE Rev. 1, 018005–018048 (2010).

47. Memmolo, P. et al. Recent advances in holographic 3d particle tracking. Adv. Opt. Photon. 7, 713–755 (2015).

48. Kamilov, U. S. et al. Optical Tomographic Image Reconstruction Based on Beam Propagation and Sparse Regularization. IEEE Trans Comput Imaging 2(1), 59–70 (2016).

49. Schürmann, M., Scholze, J., Müller, P., Guck, J. & Chan, C. J. Cell nuclei have lower refractive index and mass density than cytoplasm. J. Biophotonics 9(10), 1068–1076 (2016).

50. Yurkin, M. A. How a phase image of a cell with nucleus refractive index smaller than that of the cytoplasm should look like? J. Biophoton. 11, e201800033 (2018).

