## Supplementary Information for "Stain-free nucleus identification in holographic learning flow cyto-tomography"

### CSSI algorithm

In order to describe the steps of the proposed CSSI algorithm, sketched in Fig. 3c, we exploit the 3D numerical cell phantom shown in Figs. 3a,b.

1. The 3D RI tomogram of the analyzed cell is centered in its $L_x \times L_y \times L_z$ array, that is then divided into distinct cubes, each of which has an edge measuring $\varepsilon$ pixel, as shown in the central *xz*-slice in Fig. S1a.

The $\varepsilon$ parameter is the resolution factor at which the 3D array is firstly analyzed. It must be an even number and, after dividing each side of the 3D array by $\varepsilon$, an odd number must be obtained. Therefore, each distinct cube contains $\varepsilon^3$ voxels (i.e. RI values). The cubes completely contained within the cell shell are the investigated cubes $C_I$ (yellow cubes within the blue cell shell in Fig. S1a). The central cube is not an investigated cube, since it is taken as a reference cube $C_R$ (green cube in Fig. S1a), which vertices have *x*-, *y*-, and *z*-coordinates taken from pairs $(V_{1x}, V_{2x})$, $\left(V_{1y}, V_{2y}\right)$, and $(V_{1z}, V_{2z})$, respectively. In Figure S1b, we also display the 3D array from which the central *xz*-slice of Fig. S1a has been selected.

As discussed in the Main Text, the CSSI algorithm is based on the WMW test[32,33]. It is a rank-based non-parametric statistical test, thus distributions do not have to be normal. With a certain significance level γ, it allows rejecting or not the null hypothesis $H_0$ for which two sets of values have been drawn from the same distribution. The significance level γ is the probability of making an error of 1st species, i.e. of rejecting the null hypothesis $H_0$ when it is true. The confidence level is defined as 1-γ, i.e. it is the probability of not rejecting the null hypothesis $H_0$ when it is true. An important parameter in a statistical test is the p-value, which ranges from 0 to 1. The p-value is the observed significance level, i.e. the smallest significance level at which $H_0$ is rejected. It can be also defined as the probability of obtaining results at least as extreme as the results actually observed, when the null hypothesis $H_0$ is true. Therefore, a low p-value leads to reject the null hypothesis $H_0$, because it means that such an extreme observed result is very unlikely when the null hypothesis $H_0$ is true. In fact, if the p-value≥γ, $H_0$ is not rejected with significance level γ, while if p-value<γ, $H_0$ is rejected with significance level γ. Therefore, the greater the p-value the greater the confidence level with which two sets of values have been extracted from the same population. Hence, our algorithm performs multiple comparisons between the investigated cubes $C_I$ and the reference one $C_R$ through the WMW test, because we are assuming that the $C_R$ voxels belong to the subcellular structure of interest, thus if a certain $C_I$ has been drawn from its same distribution, then also the $C_I$ voxels belong to the subcellular structure of interest. Without loss of generality, here we are describing and testing the method in the case of the nucleus segmentation. As discussed in the Main Text, for many kinds of suspended cells (e.g., cancer cell lines) the central voxels of the cell belong to the nucleus, therefore we associate the reference cube $C_R$ to the central cube of the 3D RI tomogram.

2. An adaptive threshold $T_P$ is set according to the p-values computed through the WMW test between the investigated cubes $C_I$ and the reference cube $C_R$. It is chosen as the maximum value less than or equal to $\tau$, such that for at least one $C_I$ it happens that p-value$\geq T_P$.
3. A first rough clustering is performed through repeated $M$-iterations loops, to create a preliminary nucleus set $\mathcal{N}^P$. For each of them
    a. A temporary set $\mathcal{N}^T$ is created with the RIs of the sole reference cube $C_R$.
    b. At each of $M$ iterations
        i. A reference set $\mathcal{R}$ is created by randomly drawing $\varepsilon^3$ values from the temporary set $\mathcal{N}^T$.
        ii. For each investigated cube $C_I$, the corresponding p-value is computed with respect to the reference set $\mathcal{R}$ through the WMW test.
        iii. The investigated cubes $C_I$ such that their p-value$\geq T_P$ are added to the temporary set $\mathcal{N}^T$.
    c. After an $M$-iterations loop, all the investigated cubes $C_I$ added to the temporary set $\mathcal{N}^T$ are moved to the preliminary nucleus set $\mathcal{N}^P$, and then the temporary set $\mathcal{N}^T$ is reset.
    d. Steps a-c are repeated until at least $n$ investigated cubes $C_I$ have been stored within the preliminary nucleus set $\mathcal{N}^P$, which is shown in Fig. S1c.
4. A filtering operation is performed to delete outlier cubes from the preliminary nucleus set $\mathcal{N}^P$, thus creating a filtered nucleus set $\mathcal{N}^F$. Let $C_{\mathcal{N}^P,i}$ be a cube within $\mathcal{N}^P$, with $i = 1,2, \dots, n$.
    a. The reduced nucleus set $\mathcal{N}_i^{P-}$ is created after removing the cube $C_{\mathcal{N}^P,i}$ from the preliminary nucleus set $\mathcal{N}^P$, with $i = 1, 2, \dots, n$.
    b. A p-value vector $\bar{p}$ of length $n$ is created, which $i$-th element is the p-value computed through the WMW test between the cube $C_{\mathcal{N}^P,i}$ and the reduced nucleus set $\mathcal{N}_i^{P-}$.
    c. A distance vector $\bar{d}$ of length $n$ is created, which $i$-th element is the Euclidean distance between the centre of cube $C_{\mathcal{N}^P,i}$ and point $B$, i.e., the centroid of the preliminary nucleus set $\mathcal{N}^P$.
    d. The p-value vector $\bar{p}$ is sorted in ascending order, thus obtaining the sorted p-value vector $\bar{p}^S$, shown in Fig. S1d.
    e. The distance vector $\bar{d}$ is sorted in ascending order and normalized to its maximum, thus obtaining the sorted distance vector $\bar{d}^S$, shown in Fig. S1e by blue dots.

Both vectors $\bar{p}^S$ and $\bar{d}^S$ are used to remove outlier cubes within the preliminary nucleus set $\mathcal{N}^P$. In fact, a cube $C_{\mathcal{N}^P,i}$ is considered an outlier if it is far from the centroid $B$ and has a low p-value with respect to the other cubes in $\mathcal{N}^P$.

f. A fourth-degree polynomial is fitted to the sorted distance vector $\bar{d}^S$, thus obtaining the vector $\bar{d}^{SF}$ and its first difference $\bar{D}^{SF}$, that are reported in red in Figs. S1e,f, respectively.

g. Let $m$ be the index of the lowest value with null slope in vector $\bar{D}^{SF}$, as highlighted by the black dot in Fig. S1f. If in the vector $\bar{D}^{SF}$ there is no point with null slope, $m$ is chosen as the index of the global minimum.

h. After computing thresholds $T_1$, $T_2$, $T_3$, $T_4$, and $T_5$, the filtered nucleus set $\mathcal{N}^F$ reported in Fig. S1g is formed by cubes $C_{\mathcal{N}^P,i}$ that satisfy one of the following conditions

$$\begin{aligned} &1)\ \frac{d_i^{SF}}{d_{max}^{SF}} \leq T_1 \\ &2)\ T_1 < \frac{d_i^{SF}}{d_{max}^{SF}} \leq T_2\ \&\ p_i^S > T_4\ , \\ &3)\ T_2 < \frac{d_i^{SF}}{d_{max}^{SF}} \leq T_3\ \&\ p_i^S > T_5 \end{aligned} \tag{S1}$$

where & is the logical *and* operator, $d_i^{SF}$ and $p_i^S$ are elements of vectors $\bar{d}^{SF}$ and $\bar{p}^S$, respectively, with $i = 1,2,\dots,n,$ and $d_{max}^{SF}$ is the maximum value of vector $\bar{d}^{SF}$.

However, to build a filtered nucleus set $\mathcal{N}^F$, a strong spatial and statistical filtering has been made, in order to store only cubes that belong to the nucleus with high probability, thus leading to a strong underestimation of the nucleus region. Moreover, to increase the statistical power of the WMW test, the resolution factor $\varepsilon$ should not be too small. As a consequence, the $\varepsilon$-cubic structuring element leads to a low spatial resolution.

5. A refinement step is performed, in order to transform the filtered nucleus set $\mathcal{N}^F$ into a refined nucleus set $\mathcal{N}^R$, shown in Fig. S1h. For each cube $C_{\mathcal{N}^F,i}$ within the filtered nucleus set $\mathcal{N}^F$,
   a. Let the augmented cube $C_{\mathcal{N}^F,i}^A$ be the smallest cube centered in $C_{\mathcal{N}^F,i}$ with an edge multiple of $\varepsilon$ px, such that the p-value computed through the WMW test between its RIs and all the $\mathcal{N}^F$ values is greater than or equal to $\beta\mu(\bar{p})$, where $\mu(\cdot)$ is the average operator.
   b. To enhance the resolution, the augmented cube $C_{\mathcal{N}^F,i}^A$ is in turn divided into distinct sub-cubes with edges measuring $\varepsilon/2$ px.
   c. For each of these sub-cubes
      i. Its $\varepsilon^3/8$ values are compared with $\varepsilon^3/8$ RIs randomly drawn from the filtered nucleus set $\mathcal{N}^F$.
      ii. If the computed p-value≥ $\alpha T_P$, the examined sub-cube is inserted into the refined nucleus set $\mathcal{N}^R$.
6. All the possible pairs of sub-cubes in the refined nucleus set $\mathcal{N}^R$ are linked through a line segment.

7. A morphological closing is performed to smooth the corners of the resulting 3D polygonal and fill its holes, thus finally obtaining the partial nucleus set $\mathcal{N}_j$, displayed in Fig. S1i.
8. Steps 1-7 are repeated $K$ times on the same cell, thus obtaining $K$ partial nucleus sets $\mathcal{N}_j$, with $j = 1, 2, \dots, K$, that are slightly different from each other, because in some of the performed WMW tests, the reference set is randomly drawn from a greater one.
9. The sum of all the $K$ partial nucleus sets $\mathcal{N}_j$ provides a tomogram of occurrences, in which each voxel can take integer values $k \in [0, K]$ since each voxel may have been classified nucleus $k$ times. In Figure S2a, the central slice of this tomogram of occurrences is reported.
10. An adaptive threshold $k^*$ is set to segment the tomogram of occurrences. Let $V_k$ be the number of voxels that have been classified nucleus at least $k$ times, with $k = 1, 2, \dots, K$. Therefore, $V_1$ is the number of voxels of logical *or* among all the $K$ partial nucleus sets $\mathcal{N}_j$, while $V_K$ is the number of voxels of logical *and* among all the $K$ partial nucleus sets $\mathcal{N}_j$.
    a. A vector $\bar{V}^P$ of percentage volumes is created, which elements are computed as
    $$V_k^P = \frac{V_k}{V_1}, \tag{S2}$$
    with $k = 1,2, \dots, K$, as reported in Fig. S2b by blue dots.

    The 3D segmented nucleus-like region should be computed as the set of voxels that have occurred at least $k^{opt}$ times. The parameter $k^{opt}$ should maximize simultaneously the accuracy, sensitivity, and specificity of the proposed CSSI method. In Figure S2c, these performances are reported for each 3D segmented region composed by voxels that have occurred at least $k$ times, with $k = 1,2, \dots, K$, along with the $k^{opt}$ value, highlighted by the vertical green line. However, in a real experiment these trends are unknown, thus the intersection point cannot be computed. Therefore, a criterion is requested to find $k^*$ threshold, i.e., a suitable estimate of the $k^{opt}$ threshold.
    b. The $k^*$ threshold (red vertical line in Fig. S2b) is found as the $k$ index at which the percentage volume vector $\bar{V}^P$ is nearest to a threshold $T_V$ (orange horizontal line in Fig. S2b).

    In Figure S2c, where the $k^*$ threshold is highlighted by the red vertical line, it is clear that, despite $k^* \neq k^{opt}$, $k^*$ is located in the same quasi-constant region of $k^{opt}$, hence this estimated threshold leads to very little differences in terms of clustering performances with respect to the optimal one.
11. The final 3D nucleus-like region $\mathcal{N}$ is made of voxels that have been classified nucleus at least $k^*$ times, as shown in Fig. 3d.

In Figure 3d, it is evident that the proposed CSSI algorithm allows segmenting a 3D nucleus-like region very close to the original one, as underlined by accuracy, sensitivity, and specificity reported below the tomograms. Moreover, these values are very close to the optimal ones that could be obtained by using the optimal threshold

$k^{opt}$ instead of the estimated one $k^{*}$, i.e. $ACC^{opt} = 98.22$ %, $SENS^{opt} = 98.57$ %, and $SPEC^{opt} = 98.10$ %.

All the parameters involved in the proposed CSSI algorithm are described in Table S1. It is worth underlining that, in our experiments, a resolution factor $\varepsilon$=10 px has been set to analyse arrays made of at least $190 \times 190 \times 190$ voxels, since it was an optimum compromise between the need of having both high resolution in nucleus segmentation and high statistical power in WMW test. Anyway, in the case of tomograms with lower resolution, it can also be reduced, and all the other parameters change accordingly. However, a resolution factor $\varepsilon$ greater than 5 px is suggested in order to avoid a low statistical power in WMW test.

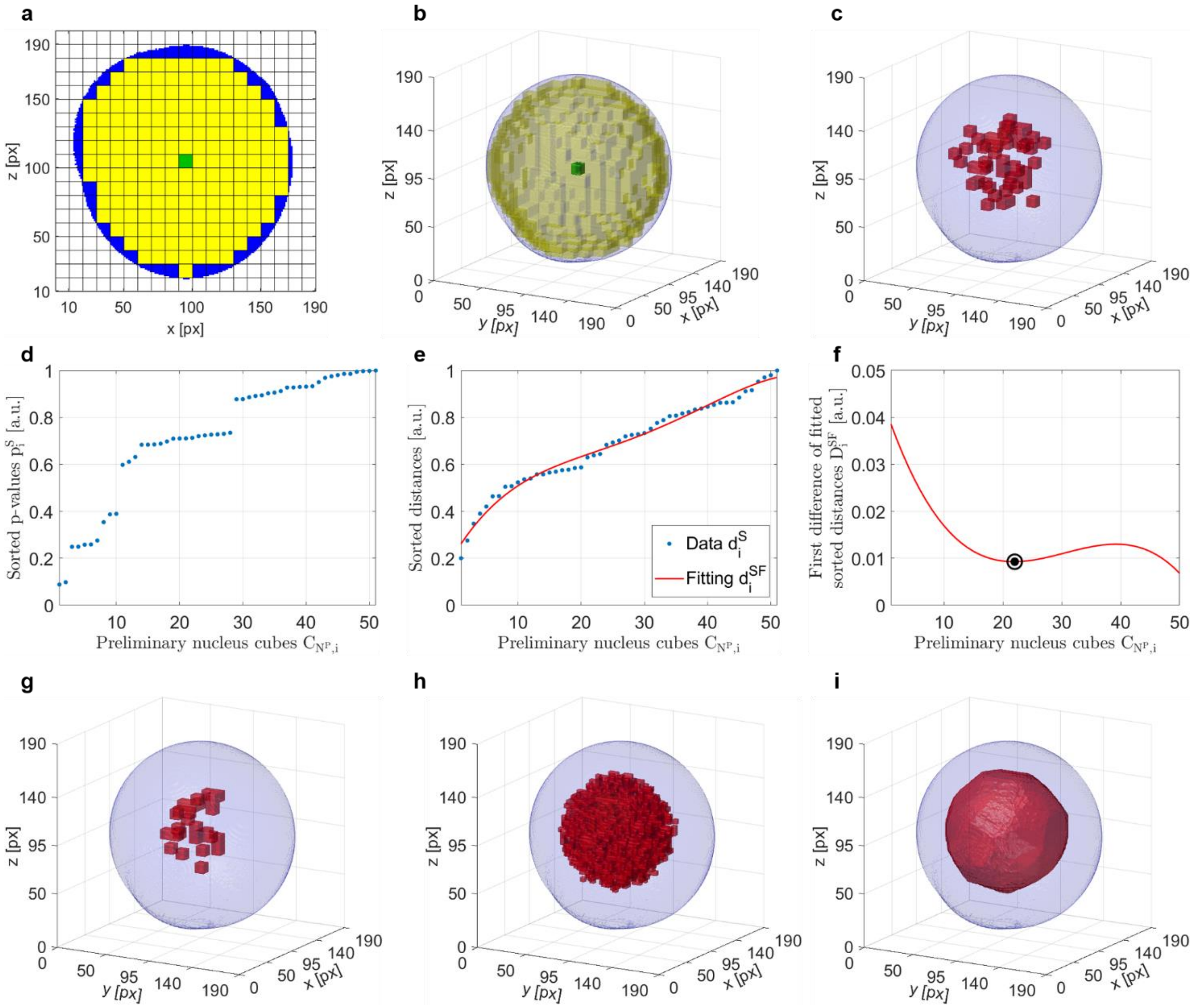


**Fig. S1. CSSI of the stain-free nucleus-like region in a 3D numerical cell phantom (Supplementary Movie 1). a** Central *xz*-slice of the 3D array divided into distinct cubes with edge $\boldsymbol{\varepsilon}$ =10 px. The central reference cube $\boldsymbol{C_R}$ is highlighted in green while the investigated cubes $\boldsymbol{C_I}$ are highlighted in yellow within the cell shell in blue. **b** 3D array from which the central *xz*-slice in (a) has been selected. **c** Preliminary nucleus set $\boldsymbol{\mathcal{N}^P}$ made of $\boldsymbol{\varepsilon}$-cubes classified nucleus after a first rough clustering. **d** Vector of sorted p-values $\boldsymbol{\bar{p}^S}$ computed through the WMW test between each cube $\boldsymbol{C_{\mathcal{N}^P,i}}$ in the preliminary nucleus set $\boldsymbol{\mathcal{N}^P}$ and the reduced nucleus set $\boldsymbol{\mathcal{N}_i^{P-}}$, with $\boldsymbol{i = 1, 2, \dots, n}$. **e** Vector of sorted and normalized Euclidean distances $\boldsymbol{\bar{d}^S}$ (blue dots) between each cube $\boldsymbol{C_{\mathcal{N}^P,i}}$ in the preliminary nucleus set $\boldsymbol{\mathcal{N}^P}$ and the centroid of all cubes in $\mathcal{N}^P$, along with the fourth-degree polynomial fitting $\bar{d}^{SF}$ (red line), with $i = 1, 2, \ldots, n$. **f** First difference $\bar{D}^{SF}$ (red line) of vector of fitted sorted distances $\bar{d}^{SF}$ in (b), with highlighted in black the lowest value with null slope. **g** Filtered nucleus set $\mathcal{N}^F$ made of $\varepsilon$-cubes classified nucleus after a spatial and statistical filtering of the preliminary nucleus set $\mathcal{N}^P$ in (c). **h** Refined nucleus set $\mathcal{N}^R$ made of $\varepsilon/2$-cubes classified nucleus after increasing resolution in the filtered nucleus set $\mathcal{N}^F$ in (d) through sub-cubes of size $\varepsilon/2$. **i** Partial nucleus set $\mathcal{N}_j$ obtained by linking sub-cubes in (e) through segment lines and by using morphological closing. In (b,c,g-i), the blue region is the cell shell and the red region is the segmented nucleus at different steps of the CSSI algorithm.

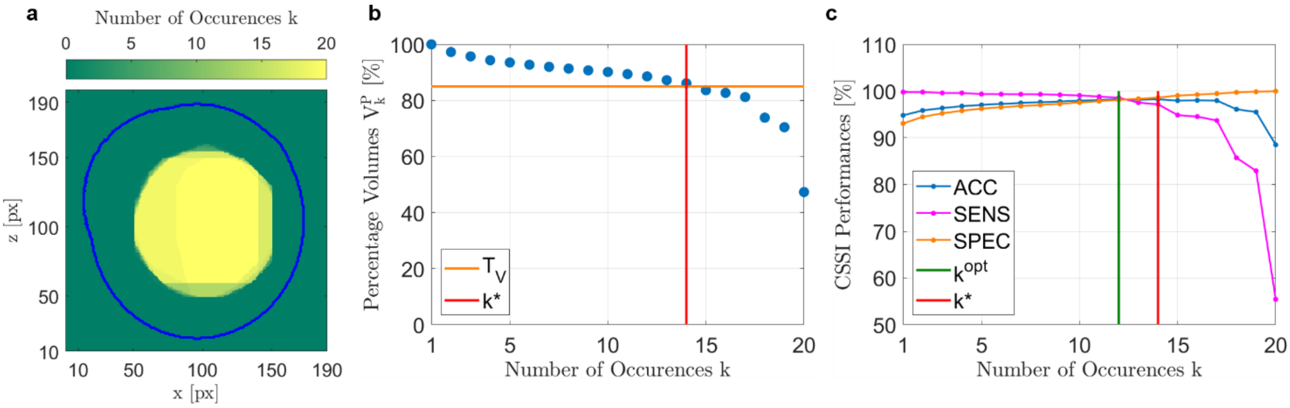


**Fig. S2. Setting of the estimated threshold $k^*$. a** Central *xz*-slice of the tomogram of occurrences, in which each voxel can take an integer value $k$ from 0 to $K = 20$, i.e. the number of times it has been classified nucleus after repeating $K$ times steps 1-7 of the CSSI algorithm on the same cell. The blue line is the cell contour. **b** Percentage volumes $V_k^P$ (blue dots), i.e. number of voxels $V_k$ classified nucleus at least $k$ times normalized to $V_{20}$, along with the threshold $T_V$ (horizontal orange line) used to find the estimated threshold $k^*$ (vertical red line) by an intersection analysis. **c** CSSI performances associated to each possible threshold $k$ for creating the final 3D nucleus-like set $N$, expressed in terms of accuracy (blue), sensitivity (magenta), and specificity (orange), along with the optimum threshold $k^{opt}$ (vertical green line in which performances are simultaneously maximized) and its estimation $k^*$ (vertical red line) computed in (b).

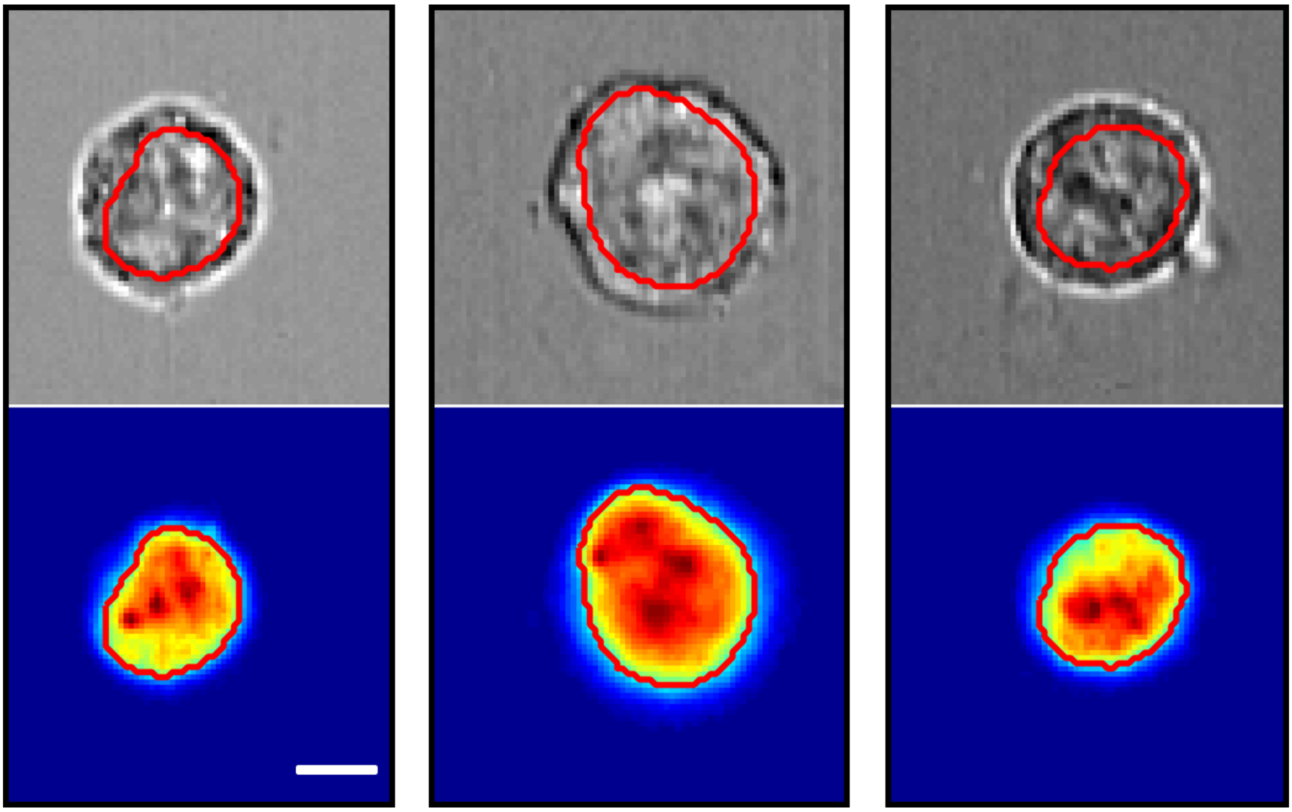

**Fig. S3. 2D cyto-fluorimetric images of SK-N-SH cells recorded by Amnis ImageStreamX®.** Three cells recorded simultaneously in brightfield images (top) and fluorescent images with the stained nucleus (bottom). The contour of the nucleus segmented by using the fluorescence information is overlapped in red. Scale bar is 5 μm.

<table>
<tr><td colspan="20">Table S1. Setting of the parameters involved in the CSSI algorithm to segment the stain-free 3D nucleus-like region from in-flow ODT reconstructions ($\lfloor\cdot\rfloor$, $\lceil\cdot\rceil$, and $\lfloor\cdot\rceil$ are the floor, ceil, and nearest integer operators, respectively).</td></tr>
<tr><td colspan="5">$\varepsilon = 10\,px$</td><td colspan="5">$V_{1x} = \frac{L_x - \varepsilon + 2}{2}$</td><td colspan="5">$V_{2x} = \frac{L_x + \varepsilon}{2}$</td><td colspan="5">$V_{1y} = \frac{L_y - \varepsilon + 2}{2}$</td></tr>
<tr><td colspan="5">$V_{2y} = \frac{L_y + \varepsilon}{2}$</td><td colspan="5">$V_{1z} = \frac{L_z - \varepsilon + 2}{2}$</td><td colspan="5">$V_{2z} = \frac{L_z + \varepsilon}{2}$</td><td colspan="5">$\tau = 0.99$</td></tr>
<tr><td colspan="5">$M = 10$</td><td colspan="5">$n = \left\lfloor \frac{2}{\varepsilon} \frac{L_x + L_y + L_z}{3} \right\rceil$</td><td colspan="5">$T_5 = \frac{1}{2}[\mu(\overline{p}) + p_{max}]$</td><td colspan="5">$T_4 = \mu(\overline{p})$</td></tr>
<tr><td colspan="10">$T_3 = \begin{cases} 0.7 & if\ \frac{m+1}{n} < 0.15 \\ 0.95 & if\ \frac{m+1}{n} > 0.85 \\ 0.7 + \frac{5}{14}\left(\frac{m+1}{n} - 0.15\right) & otherwise \end{cases}$</td><td colspan="10">$T_V = \begin{cases} \frac{\lfloor 5\mu(\overline{V}^P) \rfloor}{5} & if\ \mu(\overline{V}^P) > 50\,\% \\ \frac{\lceil 5\mu(\overline{V}^P) \rceil}{5} & if\ \mu(\overline{V}^P) < 50\,\% \\ \mu(\overline{V}^P) & if\ \mu(\overline{V}^P) = 50\,\% \end{cases}$</td></tr>
<tr><td colspan="4">$T_2 = T_3 - 0.1$</td><td colspan="4">$T_1 = T_2 - 0.1$</td><td colspan="4">$\alpha = 0.9$</td><td colspan="4">$\beta = 0.5$</td><td colspan="4">$K = 20$</td></tr>
</table>

**Table S2.** Parameters used for simulating the 3D numerical cell phantoms.

| | | | | |
|---|---|---|---|---|
| $\mu_{\mu_1} = 1.352$ | $\mu_{\mu_3} = 1.368$ | $\sigma = 0.005$ | $a_1 = 10$ | $a_2 = 1$ |
| $\mu_{\mu_2} = 1.365$ | $\mu_{\mu_4} = 1.370$ | $\sigma_\mu = 0.003$ | $b_1 = 20$ | $b_2 = 3$ |

**Table S3.** 2D morphological parameters of SK-N-SH cells measured in labelled nuclei segmented from 1820 2D FM cyto-fluorimetric images and in unlabeled nuclei segmented by CSSI algorithm from 90 2D reprojections of five in-flow ODT reconstructions.

| | | *Mean Value* | | *p-value* |
|---|---|---|---|---|
| | | *ODT* | *FM* | *WMW test* |
| *nucleus size* | *nucleus-cell area ratio [a.u.]* | 0.404 | 0.412 | 0.980 |
| *nucleus shape* | *nucleus aspect ratio [a.u.]* | 0.939 | 0.938 | 0.917 |
| *nucleus position* | *normalized nucleus-cell centroid distance [a.u.]* | 0.116 | 0.123 | 0.841 |

| **Table S4.** 3D morphological parameters of MCF-7 cells measured in labeled nuclei segmented from FM confocal images[34] and in unlabeled nuclei segmented by CSSI algorithm from three in-flow ODT reconstructions. | | | | | |
|---|---|---|---|---|---|
| | | *FM* | *ODT cell 1* | *ODT cell 2* | *ODT cell 3* |
| *nucleus size* | *nucleus-cell volume ratio [a.u.]* | 0.3396 ± 0.0939 | 0.296 | 0.415 | 0.315 |
| *nucleus shape* | *nucleus surface-volume ratio* [$\mu m^{-1}$] | 0.713 ± 0.103 | 0.702 | 0.531 | 0.635 |
| *nucleus position* | *normalized nucleus-cell centroid distance [a.u.]* | 0.152 ± 0.108 | 0.053 | 0.141 | 0.172 |
